# Cell type-specific isolation and transcriptomic profiling informs glial pathology in human temporal lobe epilepsy

**DOI:** 10.1101/2020.12.11.421370

**Authors:** Jessica Tome-Garcia, German Nudelman, Zarmeen Mussa, Elodia Caballero, Yan Jiang, Kristin G. Beaumont, Ying-Chih Wang, Robert Sebra, Schahram Akbarian, Dalila Pinto, Elena Zaslavsky, Nadejda M. Tsankova

## Abstract

The pathophysiology of epilepsy underlies complex network dysfunction, the cell-type-specific contributions of which remain poorly defined in human disease. In this study, we developed a strategy that simultaneously isolates neuronal, astrocyte and oligodendroglial progenitor (OPC)-enriched nuclei from human fresh-frozen neocortex and applied it to characterize the distinct transcriptome of each cell type in temporal lobe epilepsy (TLE) surgical samples. Differential RNA-seq analysis revealed several dysregulated pathways in neurons, OPCs, and astrocytes, and disclosed an immature phenotype switch in TLE astrocytes. An independent single cell RNA-seq TLE dataset uncovered a hybrid population of cells aberrantly co-expressing canonical astrocyte and OPC-like progenitor markers (GFAP+OLIG2+ glia), which we corroborated in-situ in human TLE samples, and further demonstrated their emergence after chronic seizure injury in a mouse model of status epilepticus. In line with their immature signature, a subset of human TLE glia were also abnormally proliferative, both in-vivo and in-vitro. Generally, this analysis validates the utility of the proposed cell type-specific isolation strategy to study glia-specific changes ex vivo using fresh-frozen human samples, and specifically, it delineates an aberrant glial phenotype in human TLE specimens.

## INTRODUCTION

Epilepsy is a debilitating neurological disorder that affects ~0.5-1% of the population^1^. The disease has been predominantly studied in the context of neuronal excitability and network dysfunction; yet, therapeutic reduction of neuronal activity has shown only limited clinical efficacy^2,3^. More recently, pathogenic roles for glia and neuroinflammation have emerged, implicating a more complex but intimate dysregulation of the glial-neuronal homeostasis in epilepsy^4,5^. Rodent studies have begun to elucidate the cell type-specific molecular pathways dysregulated in epilepsy^6^. Oligodendroglial progenitors (OPCs), also known as NG2-positive glia, have been shown to increase in number and migrate to the site of brain injury under various central nervous system (CNS) insults^7–12^, including epileptic activity^13^, where their presence has been implicated in subtle myelin dysregulation^14,15^. In contrast, astrocytes remain within their niche, where they can alter their phenotype in response to injury^16^, including in the context of seizures^17,18^. At the cellular level, reactive astrocytes in epileptic lesions show dysregulation of potassium (K+) channels, glutamate transporters, aquaporins, and connexins^6,19,20^.

Characterizing the functional and molecular biology of glia in *human* TLE pathology, however, has been much more limited, in part due to the difficulty of dissociating glia and neurons in primary tissue, the cytoplasmic processes of which are heavily interconnected^21^. Fluorescence-activated nuclei sorting (FANS) has emerged as a powerful tool to isolate and study human neuronal nuclei (NEUN+) populations from fresh-frozen archival tissue^22,23^, circumventing cytoplasmic dissociation and minimizing transcriptional activation during processing, and several recent studies have successfully profiled the full transcriptome and open chromatin landscape of neuronal NEUN+ populations in both healthy and diseases conditions^24–26^. However, similar methods to isolate specific glial subpopulations from the non-neuronal (NEUN–) element (composed of endothelium, pericytes, smooth muscle cells, inflammatory cells, and all glial subtypes) are lacking in the field and therefore much less is understood about the specific molecular alterations of glial subtypes within the diseased tissue niche^21,22,27^.

Here, we developed and validated a strategy that uses three transcription factors, NEUN, OLIG2, and PAX6, to simultaneously isolate neuronal, OPC, and astrocyte nuclei populations from non-diseased fresh-frozen postmortem human brain tissue, and then employed it, in combination with single cell RNA-seq, to characterize the cell type-specific transcriptome alterations in primary TLE neocortex.

## RESULTS

### Simultaneous isolation of astrocyte, neuronal, and OPC-enriched nuclei from bulk fresh-frozen human cortex

The role of human glia in many neurological disorders is still poorly understood due to the lack of tools that reliably isolate specific glial subpopulations from bulk tissue, directly from their native niche. To better understand the contributions of glial pathology in human epilepsy, we sought to develop a method that isolates astrocyte and oligodendroglial-lineage populations, two functionally distinct glial cell types, directly from human brain tissue. To do this, we modified the FANS NEUN+/– method for isolating neuronal nuclei from fresh-frozen cortex^22,23^ by incorporating positive selection nuclear markers for astrocyte and OPC nuclei. For OPC isolation, we used OLIG2, a known marker of adult oligodendroglial lineage cells, which shows stronger expression in OPCs compared to mature oligodendrocytes^28–30^. To find a suitable nuclear astrocytic marker, we searched for astrocyte-enriched transcription factor (TF) genes within the HepaCAM-purified resting human astrocyte transcriptome database ^31^ and found *PAX6* and *SOX9* to be among the top upregulated astrocyte-specific genes. Although largely studied in the context of early neuroepithelial development and retinal neuronal specification^32–34^, PAX6 has been shown to promote the maturation of murine astrocytes^35^ as well as to co-express with the astrocytic marker GFAP in epileptic human tissue ^36^, and SOX9 is widely expressed by mouse and human astrocytes^27^.

We then isolated nuclei from non-diseased (control) postmortem temporal neocortex (TL) containing gray and white matter, performed FANS using the three positive selection TF markers, and detected reliable separation of NEUN+, PAX6+ / SOX9+ (NEUN–OLIG2–), and OLIG2+ (NEUN–PAX6–/SOX9–) populations, hereafter referred to as NEUN+, PAX6+ / SOX9+, and OLIG2+ for simplicity (Fig. 1a, Supplementary Fig. 1a-b). Analysis of canonical lineage-specific markers in each of these populations by RT-qPCR confirmed strong enrichment of neuronal, astrocytic, and OPC markers in the respective populations. PAX6+ nuclei were distinctly enriched for the astrocytic markers *ALDH1L1* and *GFAP*, as well as for *PAX6* (Fig. 1c). SOX9+ nuclei were also enriched for astrocytic markers, but to a slightly lesser extent (Supplementary Fig. 1c), prompting subsequent FANS experiments to be performed with PAX6 only. OLIG2+ nuclei were enriched for the OPC markers *OLIG2*, *CSPG4* (NG2) and *PDGFRA* (Fig. 1c), and, importantly, showed low expression for the myelinating oligodendrocyte marker *PLP1* (Fig. 1d). Instead, *PLP1* expression was enriched in a distinct population of nuclei derived from gating on low OLIG2 expression (OLIG2^LOW^) (Fig. 1b, 1d; Supplementary Fig. 1a). This suggested that excluding the OLIG2^LOW^ fraction of the OLIG2+ population (as shown in Fig. 1b and Supplementary Fig. 1a) can enrich for OPCs, relative to myelinating oligodendroglial populations, and this gating strategy was used for all subsequent sorting experiments. Finally, the NEUN–PAX6–OLIG2– triple negative (TN) population was depleted of astrocyte, OPC, and neuronal markers (Fig. 1c) but showed strong enrichment for the microglial marker *CD11b* (Supplementary Fig. 1d).

**Fig. 1.**
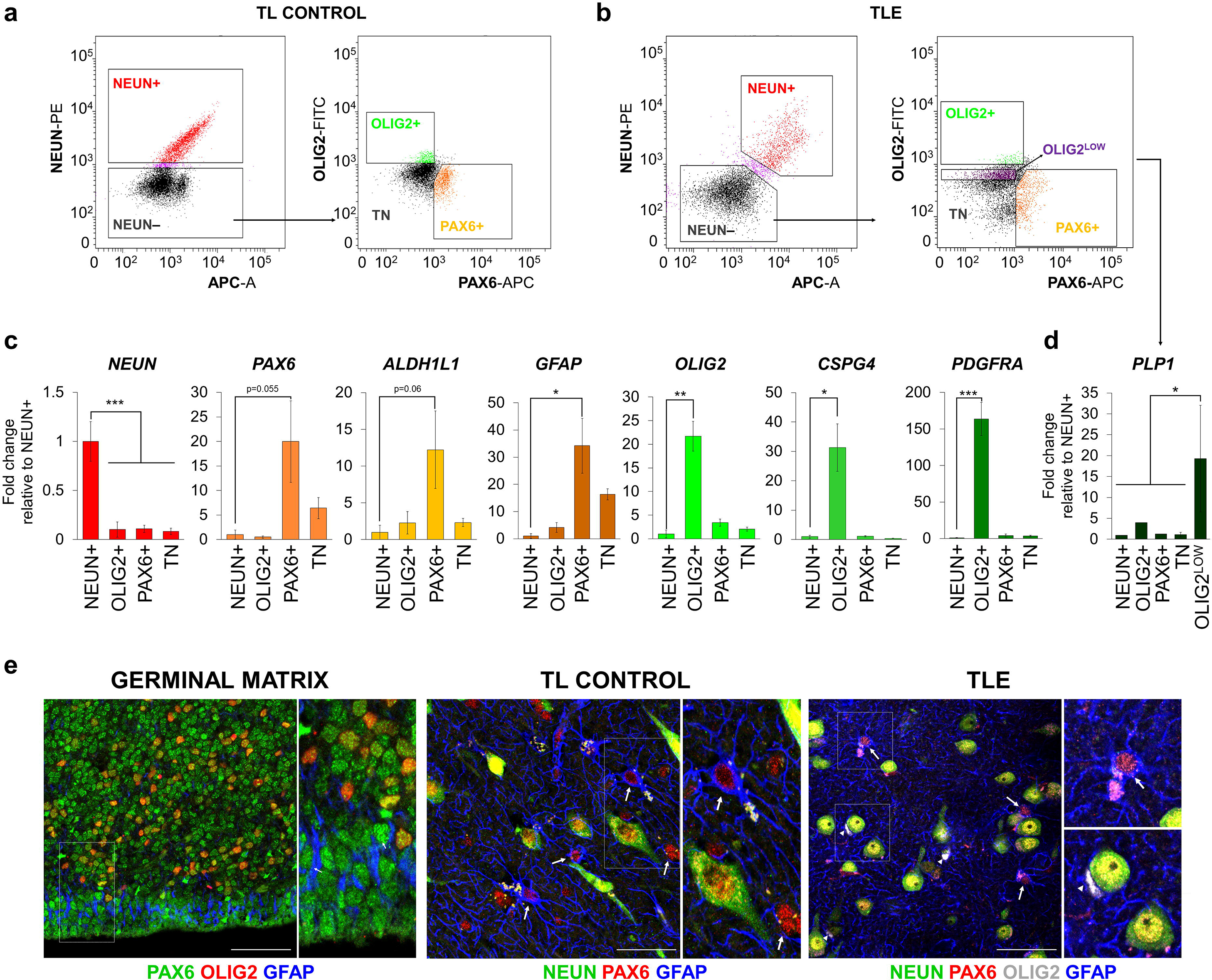
Simultaneous isolation of astrocyte, neuronal, and OPC-enriched nuclei from human temporal neocortex. **(a-b)** Fluorescence-activated nuclei sorting (FANS) using anti-NEUN, anti-PAX6, and anti-OLIG2 antibodies simultaneously isolates three distinct nuclei populations from human postmortem control (TL) and epilepsy (TLE) temporal lobe neocortex: NEUN+, PAX6+(NEUN–), and OLIG2+(NEUN–), excluding the OLIG2^LOW^ fraction. TN = triple negative (NEUN–PAX6–OLIG2–) population. See also Supplementary Fig. 1a. **(c)** Gene expression analysis by RT-qPCR confirms high expression of the genes used as markers for isolation and shows significantly enriched expression of the astrocytic markers *GFAP* and *ALDH1L1* in PAX6+( NEUN–) nuclei and of the OPC markers *CSPG4* (NG2) and *PDGFRA* in OLIG2+( NEUN–) nuclei (n = 4 TL control brains). Bars represent mean ± SEM. P-values calculated from one-tailed t-test compared to NEUN+ population. PAX6+ vs. NEUN+: *PAX6* p = 0.055; *ALDH1L1* p = 0.067; *GFAP* p = 0.019. OLIG2+ vs. NEUN+: *OLIG2* p = 0.0014; *CSPG4* p = 0.012; *PDGFRA* p = 0.0008. **(d)** Quantification of expression of the myelinating oligodendrocytic marker *PLP1* by RT-qPCR, showing its significantly higher expression in the OLIG2^LOW^ gated nuclei population compared to all others. Bars represent mean ± SEM, (n=3 TLE brains). *p<0.05 one-tailed t-test. **(e)** Representative immunofluorescence images of PAX6, OLIG2, and NEUN expression in developing germinal matrix (left), adult postmortem TL neocortex (center) and adult TLE neocortex (right). In adult TL and TLE neocortex, PAX6 expression is seen in GFAP+ astrocytes (arrows) and NEUN+ neurons but is absent in OLIG2+ oligodendroglial cells (arrowheads). Scale bar = 50μM. See also Supplementary Fig. 1e.

We also characterized the cell type-specific distribution of OLIG2, PAX6, and NEUN protein expression in situ, in adult TL and TLE neocortex, as well as in developing brain as a control. PAX6 was expressed widely during neurodevelopment in human anterior germinal matrix tissue (18-20 gestational weeks), where it co-localized with presumed GFAP+ radial-like glia and OLIG2+ glial progenitors (Fig. 1e). In adult tissues, PAX6 was expressed strongly in (GFAP+) astrocytes in both normal adult and epileptic adult temporal lobe neocortex, and its expression did not appear to overlap with OLIG2-positive cells, as seen during development (Fig. 1e). In adult tissues, NEUN was expressed exclusively in neurons and OLIG2 was expressed by PAX6– glia in both TL control (Supplementary Fig. 1e) and TLE tissues (Fig. 1e). Interestingly, while NEUN+ nuclei showed very low expression of *PAX6* (Fig. 1c), in line with previous human transcriptome studies^31^, we detected weak PAX6 immunoreactivity in adult TL neurons (Fig. 1e). This discrepancy did not affect the FANS isolation strategy, since PAX6+ astrocytes were isolated from the NEUN– fraction. Overall, the gene and protein expression patterns of NEUN, PAX6, and OLIG2 provided confidence in their use for simultaneous isolation of adult NEUN+ neuronal, PAX6+ (NEUN–) astrocyte, and OLIG2+ OPC-enriched nuclei populations.

### Nuclear RNA-seq validates FANS cell type specificity

To further validate this FANS isolation strategy, we analyzed the full nuclear transcriptome of NEUN+, PAX6+, and OLIG2+ nuclei populations isolated from non-diseased fresh-frozen human postmortem TL and from pathological TLE neocortex, both containing gray and white matter (Supplementary Table 1). We focused specifically on the less well-characterized human neocortex (rather than hippocampus) of non-lesional TLE samples, which contained diffuse subpial (Chaslin) and neocortical astrogliosis on diagnostic neuropathology, including away from sites of electrode placement (Table 1). In order to relate nuclear RNA signatures to gene expression, we made the assumption that a steady state of mature mRNA is being maintained between the nucleus and the cytoplasm by continuous RNA synthesis, transport, and turnover^37–39^. Indeed, several studies have now demonstrated that nuclear and total cellular mRNA pools and transcriptome signatures are comparable^40–42^.

**Table 1.** Sample information. PMT: Postmortem time; IF: Immunofluorescence; M: Male, F: Female; yo: years old; h: hours; TS: Tuberous sclerosis. FCD = Focal cortical dysplasia. * Hippocampal sclerosis seen away from the sampled area; ** tubers seen away from the sampled area.**✓**’ Samples included in the IF analysis of Ki67 proliferation cell type distribution.

| SAMPLE ID | TISSUE, LOCATION | Age, Gender PMT | NUCLEI RNA-SEQ | SC RNA-SEQ | PATHOLOGICAL DIAGNOSIS |
| --- | --- | --- | --- | --- | --- |
| **Cell-type specific FANS Validation, TL vs. TLE** |  |  |  |  |  |
| 10662 | TL control, temporal lobe neocortex | 34yo, M<br>12h<br>PMT | ✓ |  | No neuropathological changes seen in cortex |
| 10355 | TL control, temporal lobe neocortex | 45yo, F<br>12h<br>PMT | ✓ |  | No neuropathological changes seen in cortex |
| 10997 | TL control, temporal lobe neocortex | 27yo, M<br>21h<br>PMT | ✓ |  | No neuropathological changes seen in cortex |
| 12321 | TLE, temporal lobe neocortex | 50yo, M | ✓ |  | Cortical and Chaslin Gliosis; White matter neuronal heterotopia |
| 10308 | TLE, temporal lobe neocortex | 29yo, F | ✓ |  | Cortical and Chaslin gliosis; FCD Ia |
| 12726 | TLE, temporal lobe neocortex | 13yo, M | ✓ |  | Cortex without pathologic changes; * |
| 14431 | TLE, temporal lobe neocortex | 28yo, M |  | ✓ (10X, v2) | Cortical Gliosis; Ectopic white matter neurons with mild dysmorphic changes |
| 19619 | TLE, temporal lobe neocortex | 31yo, F |  | ✓ (10X, v2) | Chaslin gliosis; * |
| 20188 | TLE, temporal lobe neocortex | 58yo, F |  | ✓ (10X, v3) | Cortical and Chaslin gliosis, rare mildly dysplastic neurons; * |
| 12814 | TLE, temporal lobe neocortex | 11yo, M |  | ✓ (10X, v1) | Chaslin gliosis; * |
| 13059 | TLE, temporal lobe neocortex | 8yo, M |  | ✓ (10X, v1) | Gliosis, neuronal heterotopia; |
| SAMPLE ID | TISSUE, LOCATION | Age, Gender | Ki-67 IF | IN VITRO ASSAYS | PATHOLOGICAL DIAGNOSIS |
| **Phenotypic analysis of epilepsy glia** |  |  |  |  |  |
| 12321 | TLE, temporal lobe neocortex | 50yo, M | ✓' (high) | ✓ | Cortical and Chaslin Gliosis; White matter neuronal heterotopia |
| 10308 | TLE, temporal lobe neocortex | 29yo, F | ✓ (low) | ✓ | Cortical and Chaslin gliosis; FCD Ia |
| 12726 | TLE, temporal lobe neocortex | 13yo, M | ✓' (high) |  | Cortex without pathologic changes; * |
| 14431 | TLE, temporal lobe neocortex | 28yo, M | ✓ (low) | ✓ | Cortical Gliosis; Ectopic white matter neurons with mild dysmorphic changes |
| 12319 | Epilepsy, frontal lobe neocortex | 27yo, M | ✓' (high) |  | Chaslin Gliosis; Ischemic changes |
| 12433 | Epilepsy, TS, frontal lobe neocortex | 4yo, M | ✓ (low) | ✓ | Cortical and Chaslin Gliosis; ** |

We first performed unsupervised clustering and principal component analysis (PCA) of all sequenced samples, which separated according to cell type (Fig. 2a, Supplementary Fig. 2). As expected, glial populations (astrocytes and OPCs) were more similar to one another than to neuronal populations in the first PC dimension, and astrocytes separated from OPCs in the second PC dimension, in both TL and TLE tissues (Fig. 2a). Analysis of canonical lineage-specific markers corroborated the specificity of our isolation technique for astrocyte and OPC populations in both TL and TLE tissue types, despite notable downregulation of several canonical astrocyte markers in the epilepsy samples (Fig. 2b). PAX6+ nuclei populations were strongly enriched for astrocytic markers, while OLIG2+ nuclei were enriched for OPC and pan-oligodendroglial markers; and both showed minimal expression of vascular, inflammatory, and neuronal-specific markers (Fig. 2b). Using gene set enrichment analysis (GSEA)^43^, we also compared how TL control FANS nuclear transcriptomes relate to previous whole cell transcriptome data obtained from purified human astrocytes^31^ and mouse OPCs^44^. We found significant enrichment of the top HepaCAM-purified human astrocyte genes^31^ within PAX6+ transcriptomes (Fig. 2c), and of the top mouse OPC genes^44^ within OLIG2+ transcriptomes (Fig. 2c). Thus, nuclear RNA-seq defined distinct cell type-specific transcriptome signatures in the three sorted populations and corroborated the concordance between nuclear and whole cell RNA for highly expressed and cell lineage-specific transcripts. This provided confidence in using this immunotagging strategy for further analysis of cell type-specific transcriptome dysregulation in the context of epilepsy.

**Fig. 2.**
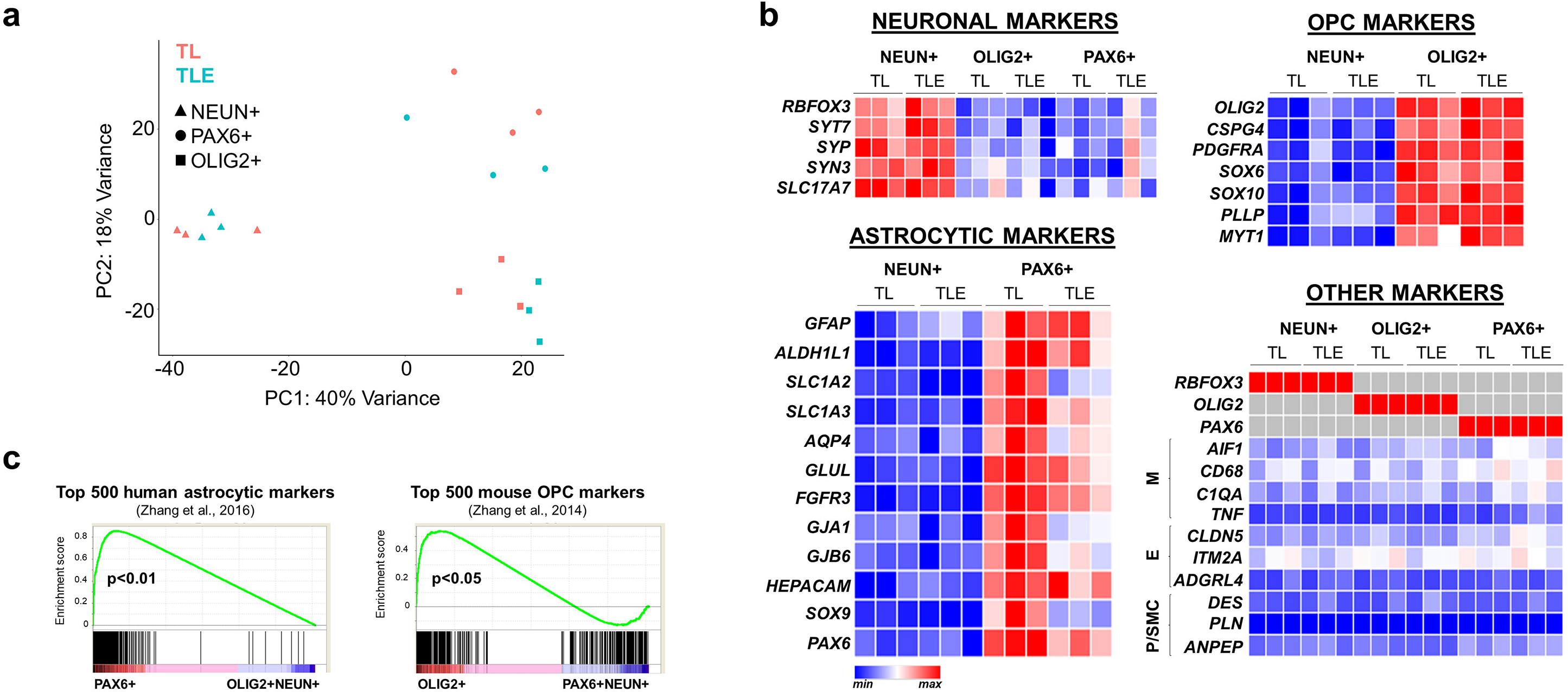
Nuclear RNA-seq confirms cell type specificity in TL and TLE FANS-isolated populations. **(a)** RNA-seq principal component analysis reveals separation driven by sorted cell type. See also Supplementary Fig. 2. Note greater similarity between TLE astrocyte and OPC transcriptomes, compared to control TL astrocyte in the PC2 dimension. **(b)** Heatmap representations of nuclear gene expression (rld-normalized) RNA-seq data, derived from sorted PAX6+, NEUN+, and OLIG2+ populations. Strong and selective enrichment of astrocytic, neuronal, and OPC markers is seen in each respective FANS population (normalized by row), with lack of contaminant microglial (M), endothelial (E), and pericyte / smooth muscle cell (P/SMC) gene expression in the sorted populations (normalized by column). **(c)** Gene set enrichment analyses (GSEA) confirm significant, cell type-specific enrichment. Rld-normalized nuclei RNA-seq data (PAX6+, OLIG2+, NEUN+) is analyzed against the top 500 overexpressed set of genes unique to resting human astrocytes ^31^ or to mouse OPCs ^44^. For each gene set, each gene expression value is calculated relative to the average expression of all other populations and then sorted by highest relative expression.

### Functional enrichment analysis of dysregulated genes in distinct human epilepsy cell types

Next, we performed a series of differential transcriptome analyses (TLE vs. TL) in order to define the epilepsy-specific and cell-type-specific dysregulated genes in TLE astrocytes, OPCs, and neurons (Supplementary Table 2), and used functional enrichment tools to characterize the most significantly enriched biological processes defined by up- and downregulated genes in each TLE cell type (Fig. 3a, Supplementary Tables 3-4). In general, we observed a net loss of gene expression in epilepsy compared to postmortem control, in each cell type population, detecting a larger number of downregulated than upregulated genes. In astrocytes (Astrocyte TLE vs. Astrocyte TL control), differentially dysregulated genes were highly enriched for the functional gene ontology (GO) terms: “nervous system development”, “gliogenesis”, “glucose metabolic process”, “response to wounding”, “regulation of extracellular matrix (ECM) organization”, “L-glutamate transmembrane transport”, “stabilization of membrane potential”, “cellular potassium ion transport”, and “DNA replication” (Fig. 3a, Supplementary Tables 3-4). Differentially dysregulated genes in OPCs (TLE vs. TL control) were enriched for GO terms related to “protein folding”, “nervous system development”, “myelination”, “chromatin maintenance”, “G2/M transition of mitotic cell cycle”, “DNA repair”, “lipid digestion”, and “tyrosine catabolic process” (Fig. 3a, Supplementary Tables 3-4). Differentially dysregulated genes in neurons (TLE vs. TL) yielded highest functional significance for GO terms related to “cell communication”, “signal transduction”, “regulation of exocytosis”, “synaptic vesicle transport”, “semaphorin-plexin signaling pathway involved in axon guidance”, “actin cytoskeleton reorganization”, “regulation of high voltage-gated calcium channel activity”, and “signal transduction” (Fig. 3a, Supplementary Tables 3-4). The majority of differentially expressed genes in these analyses were cell-unique as they were not significantly dysregulated in the other cell-type comparisons (Supplementary Table 3). Excluding the small subset of non-unique differential genes did not significantly alter the top enriched functional biological processes in each dataset analysis (Supplementary Table 3).

**Fig. 3.**
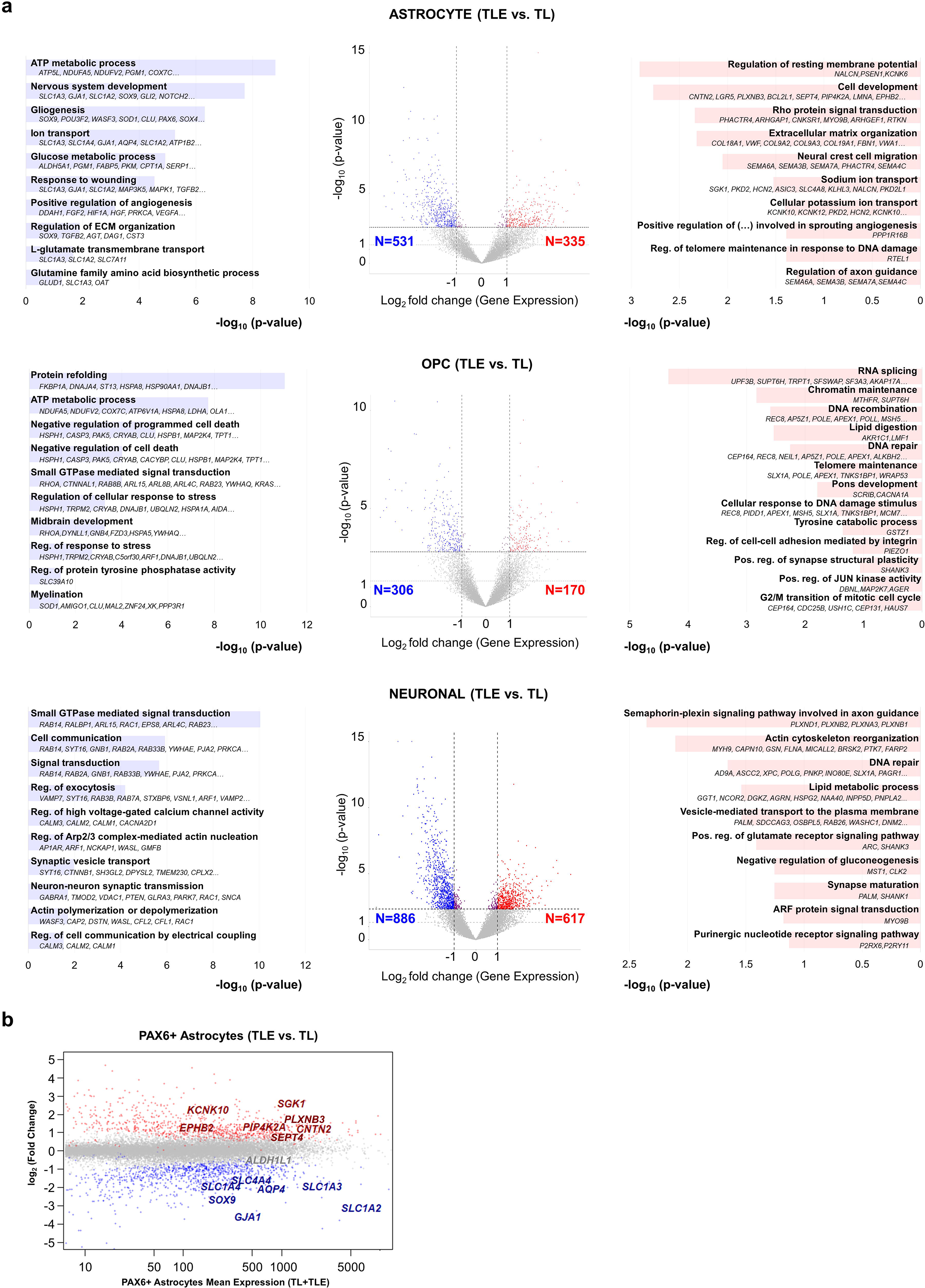
Dysregulated genes and biological processes in human temporal lobe epilepsy astrocyte, OPC and neuronal populations. **(a)** Differential expression analysis (TLE vs. TL) for each sorted cell type is represented by a volcano plot, depicting significantly upregulated (red) and downregulated (blue) genes in the epilepsy astrocyte, OPC, and neuronal populations. Functional enrichment analyses, performed using HOMER and DAVID (D) tools, depict top-enriched biological processes dysregulated in epilepsy, for each cell type (log_2_ fold change < −1 for downregulated and > 1 for upregulated, Benjamini-Hochberg adjusted p-value < 0.1). See Supplementary Tables 3 and 4 for complete list. **(b)** MA plot shows the relative expression of genes differentially up- and down-regulated in PAX6+ epilepsy astrocytes, marked by red and blue dots, respectively.

Overall, these enrichment analyses recapitulated cell-type-specific processes related to astrocyte, OPC, and neuronal function previously established in mouse models, and also uncovered several still poorly understood glial-specific pathological changes in the context of human TLE. One striking example was the significant alteration in the phenotype of epilepsy astrocytes towards de-differentiation, with downregulation of genes important for normal maintenance of synaptic homeostasis, and glial differentiation. Among the most robustly, significantly and uniquely downregulated genes in epilepsy astrocytes were the glutamate transporter GLAST/EAAT1 (*SLC1A3*); the sodium-dependent neutral amino acid transporter ASCT1 (*SLC1A4*); the gap junction proteins connexin 30 (*GJB6*) and connexin 43 (*GJA1*); and the water channel *AQP4* (Fig. 3b). GLT-1/EAAT2 (*SLC1A2*) was significantly downregulated in both TLE astrocytes and TLE neurons, with the relative expression of *SLC1A2* in astrocytes being much higher than in neurons (Supplementary Tables 1-2). In contrast, significantly and uniquely upregulated genes in TLE astrocytes included many related to ECM organization and cell development (Fig. 3b, Supplementary Tables 1-4), overall pointing towards a phenotypic switch in epilepsy astrocytes from mature synaptic maintenance to extracellular remodeling, with possible tendency towards immature de-differentiation.

### Single cell RNA-seq uncovers a subset of TLE glia co-expressing OPC and astrocyte lineage markers

To further scrutinize the cellular heterogeneity of astrocytes and OPCs in human epilepsy, we also performed single cell RNA-seq (scRNA-seq) on fresh temporal neocortex derived from five patients with medically refractory TLE, representing the primary recently electrode-mapped seizure focus. Overall, we sequenced a total number of 24,025 single cells, with an average depth of 113,788 mean reads/cell (Supplementary Table 5). The data was processed with the Cell Ranger pipeline (10X Genomics) for all five samples, and, to elucidate distinct cell types, clustered using Cell Ranger (Supplementary Fig. 3; all samples), Seurat^45^ (Fig. 4; samples 14431, 19619, and 20188) and MetaCell^46^ (Supplementary Fig. 5; sample 14431). Based on canonical cell lineage markers, we identified distinct subpopulations of all expected neural cell types, as well as an unexpected hybrid glial population with both OPC and astrocyte marker identity (Fig. 4a-c, Supplementary Fig. 3, Supplementary Fig. 4). Inflammatory and vascular cells were overrepresented in the scRNA-seq data, whereas neurons were underrepresented. Since we readily detected NEUN+ neuronal nuclei from frozen TLE tissue (Fig. 1–2), the underrepresentation of neurons in our scRNA-seq datasets collected from intact single cells could have reflected the enhanced sensitivity of neuronal membrane and cytoplasm to ischemia, and less likely loss during the droplet-based processing.

**Fig. 4.**
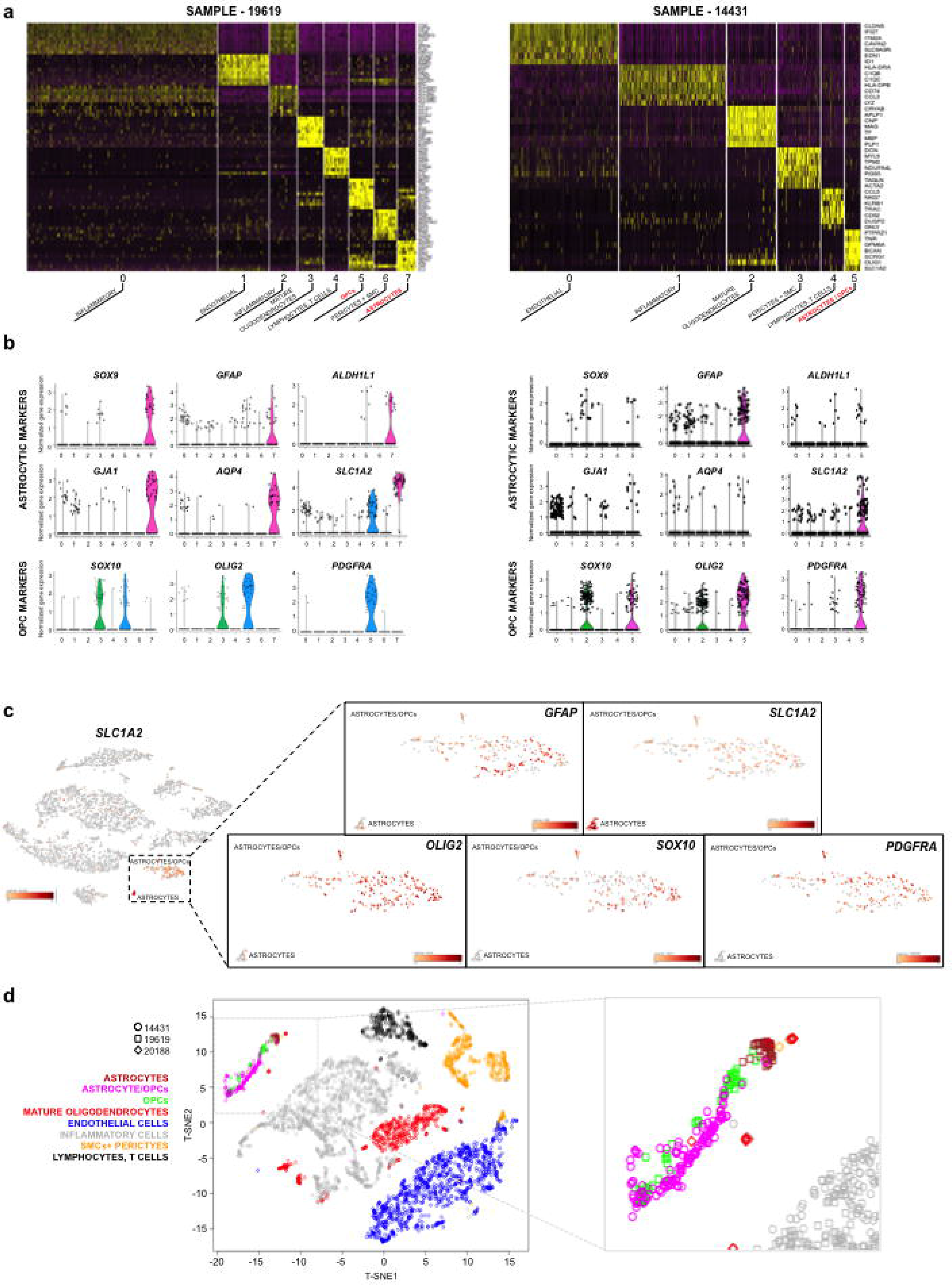
Single cell RNA-seq analysis in human TLE neocortex uncovers a hybrid astrocyte/OPC population. **(a-b)** Heatmap of Seurat-based unsupervised clustering in the two TLE samples with highest data quality, with cell-type annotation based on canonical lineage marker expression. Violin plots show cluster distribution of selected canonical astrocyte and OPC-lineage markers for each sample (b). While astrocyte and OPC markers in sample 19619 are largely mutually exclusive, annotating to respective clusters 7 and 5, sample 14431 shows a single astrocyte/OPC hybrid cluster co-expressing both cell-type markers. **(c)** T-distributed stochastic neighbor embedding (T-SNE) visualization of unsupervised clustering for sample 14431, showing a distinct small population of astrocytes expressing canonical mature astrocyte markers (*SLC1A2* shown), detected only using Cell Ranger-based clustering metrics, next to the larger astrocyte/OPC population defined by co-expression of astrocyte markers (*GFAP*, *SLC1A2* shown) and OPC markers (*OLIG2, SOX10*, and *PDGFRA* shown). Note the presence of hybrid expression of *GFAP* and *OLIG2/SOX10/PDGFRA* in a subset of cells within this cluster (zoomed in images on the right). Gene expression values are scaled for each marker (bars, bottom). **(d)** Integrative analysis of the three TLE samples processed using mnnCorrect for batch sample correction. Seurat-defined cell-type clusters are represented by cells from three independent TLE samples, including the astrocyte/OPC cluster, with variable frequency. Although Seurat lacked the resolution to identify a separate astrocyte and/or OPC cluster for sample 20188, the power of the integrative analysis allowed localization of individual 20188 cells to those clusters.

The samples with highest quality data were used for subsequent integrative analyses. We used mnnCorrect^47^ to correct for sample batch effects, which preserved the cell type-specific clustering structure and showed contribution of each sample to most clusters, albeit with variable frequency (Fig. 4d). Overall, the clustering algorithms identified at least two distinct subpopulations of astrocytes in the TLE samples (Fig. 4, Supplementary Fig. 3). The first one expressed canonical mature astrocyte markers, such as *SLC1A2*, *GJB6*, *GJA1*, *AQP4*, and *SOX*9 (Fig. 4a-b left, cluster 7; Fig. 4c; Supplementary Fig. 3), which have been previously used to cluster “normal” astrocytes in human temporal neocortex using single cell RNA-seq^48^. A second cluster was noted in all TLE samples, with variable representation in each, defined by the simultaneous expression of canonical astrocytic markers, such as *GFAP* or *SLC1A2,* along with canonical OPC markers, such as *OLIG2* or *PDGFRA* (Fig. 4a-b right, cluster 5; Fig. 4c; Supplementary Fig. 3). Careful analysis within the “astrocyte/OPC” cluster revealed many single cells with dual expression of astrocyte and OPC-lineage markers (*GFAP*+/*SLC1A2*+ and *OLIG2*+/*SOX10*+/*PDGFRA*+) (Fig. 4c, Supplementary Fig. 3), suggesting a hybrid astrocyte/OPC phenotype. We considered the possibility that dual lineage cells may represent doublet artifact, which we estimated in the 1-5% range for the 10X nanofluidic pipeline. However, the observed hybrid phenomenon was seen preferentially in the astrocyte/OPC cluster, with variable frequency among samples (Supplementary Fig. 3), and it was associated with random distribution of UMI cell counts (Supplementary Table 5), stochastically arguing against this being a consequence of artifact. As expected, differential expression and functional GO term enrichment analyses of this hybrid TLE glial cluster showed dual functional phenotypes (“astrocyte differentiation” and “oligodendrocyte differentiation”) as well as enrichment of several dysregulated pathways observed in the previous bulk-sorted nuclear RNA-seq TLE vs. TL glial analyses (Supplementary Table 3).

To try and disambiguate the “astrocyte/OPC” hybrid population, we performed additional computational analysis on the TLE sample where this population was most highly represented, sample 14431. Both Cell Ranger and Seurat were unable to distinguish astrocytes and OPCs from the seemingly hybrid “astrocyte/OPC” population (Fig. 4, Supplementary Fig. 3). MetaCell was the only clustering tool able to further split the “astrocyte/OPC” population. For control, we took advantage of C1 Fluidigm single cell RNA-seq data performed on normal TL tissue away from the seizure focus in patients with TLE^48^, to identify defining normal human temporal neocortex astrocyte and OPC markers. The “astrocyte/OPC” TLE cluster could not be annotated using “normal” astrocyte markers ^31,48^; instead, the hybrid population was annotated using “normal” OPC markers^48^ (Supplementary Fig. 5a, left). Interestingly, the only marker able to dissociate the astrocyte/OPC cluster in the MetaCell analysis was *GFAP* (Supplementary Fig. 5a right, 5b), an intermediate filament expressed in astrocytes and upregulated under various reactive conditions.

### Olig2+/Gfap+ glia emerge after pilocarpine-induced status epilepticus

To corroborate the presence of glial cells co-expressing astrocytic and OPC canonical markers as a consequence of seizures in a controlled experimental setup, and in the absence of depth electrode placement, we turned to a mouse model of epilepsy^49^ and examined the brains of mice with pilocarpine-induced status epilepticus, 7-11 weeks after their initial documented seizure activity. Immunohistochemical analysis confirmed the presence of reactive astrogliosis only in mice with pilocarpine-induced seizures, present up to 2.6 months after seizure induction, which was seen throughout the brain but was most prominent within the temporal cortex and hippocampus. Within the frontal and temporal cortex of epilepsy mice, but not in controls, we detected markedly reactive, Gfap++ cells with bonafide astrocyte morphology, a subset of which showed weak-to-strong co-expression of Olig2 (Fig. 5). We also detected many Olig2 positive oligodendroglial lineage cells, which lacked noticeable Gfap expression. This analysis confirmed the emergence of glia co-expressing Gfap and Olig2 as a consequence of epileptic seizures, which morphologically resembled reactive astrocytes.

**Fig. 5.**
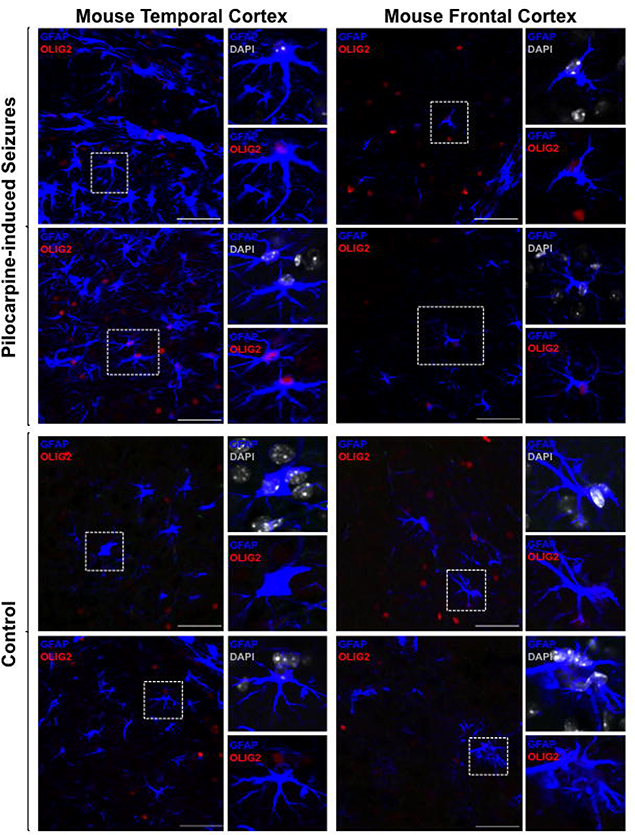
Olig2+/Gfap+ glia emerge in pilocarpine-induced seizures. Representative immunohistochemical images demonstrate the presence of Gfap++ reactive astrocytes in mice with pilocarpine-induced seizures, seven weeks after treatment, a subset of which co-express the OPC marker Olig2. Littermate-matched control mice do not display Gfap/Olig2 co-localization. Representative images from two different mice are shown for each condition.

### A subset of GFAP+ TLE glia display an aberrant proliferative phenotype

To confirm the presence of GFAP+OLIG2+ glia in human TLE samples at a protein level, we performed immunofluorescence analysis in primary TLE tissue using anti-OLIG2, and anti-GFAP antibodies. We also included anti-Ki67 antibody in the experiment to evaluate the proliferative status of this hybrid population, given its immature transcriptome signature. In half of TLE samples, we detected higher degree of cell proliferation exhibited by Ki67 immunoreactivity (Fig. 6a-c, Table 1), than expected for the largely quiescent normal TL parenchyma^50^. Within the pool of proliferative Ki67+ TLE cells, a small subset were indeed double positive for GFAP and OLIG2 (Fig. 6b-c), corroborating the presence of GFAP+OLIG2+ hybrid glia in human TLE tissue, with proliferative potential. Intriguingly, a few Ki67+ cells were GFAP+ astrocytes (Ki67+GFAP+OLIG2-) (Fig. 6b-c), which are typically quiescent under physiological conditions^50,51^. As expected, the majority of the remaining Ki67+ cells corresponded to microglia (IBA1+), and proliferative OPCs (Ki67+OLIG2+GFAP-) (Fig. 6a-c).

**Fig. 6.**
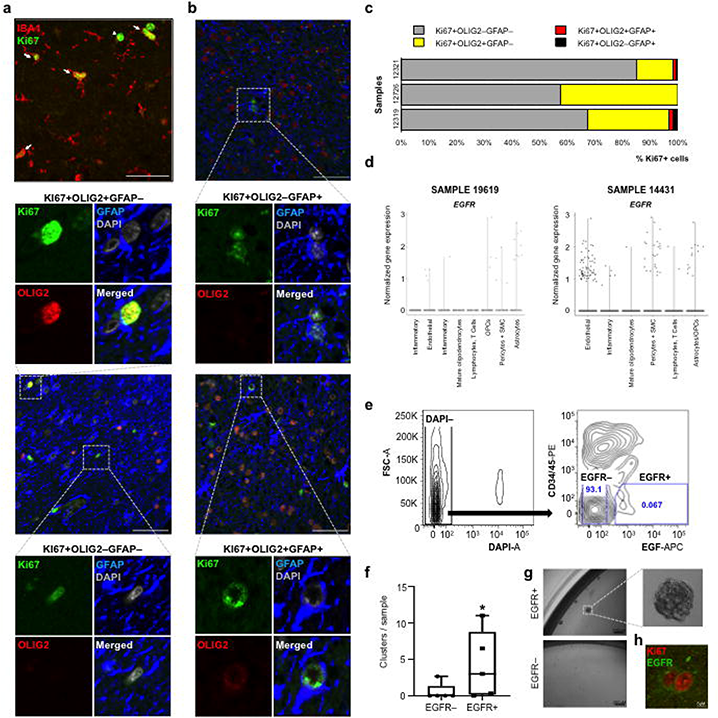
Glial proliferation in human temporal lobe epilepsy. **(a-b)** Representative immunofluorescence images from the subset of TLE samples with high proliferation, as assessed by Ki67 labeling. A large portion of Ki67-positive cells co-express the microglial marker IBA1 (AIF1) (arrows), although Ki67+IBA1**–** cells are not uncommonly seen as well (arrowheads) (a). Representative immunofluorescence images of GFAP, OLIG2 and Ki67 co-expression in TLE tissue (b). **(c)** Quantification of cell-type-specific contribution to proliferation in the subset of TLE samples with high proliferation index, assessed by co-expression of Ki67 with OLIG2 and/or GFAP. Ki67+ cells were examined within the entire TLE tissue section, 14-58 40X fields overall. **(d)** Single cell RNA-seq analysis reveals localization of EGFR+ cells to distinct cell type clusters: astrocytes, OPC/astrocytes, pericytes, inflammatory and endothelial cells (n=2 shown, observed in all samples). **(e)** FACS plots showing isolation strategy to purify EGFR+ (CD34–CD45–) cells from fresh TLE tissue. A small but distinct population of EGFR+ (DAPI–CD34– CD45–) cells are isolated, and subjected to primary cell culture along with EGFR– cells. FACS excludes dead cells (DAPI+), endothelial (CD34+) and inflammatory (CD45+) cells from the culture analysis. Numbers in boxes indicate % from total DAPI– cells. **(f)** EGFR+ cells form proliferative clusters in vitro, significantly more compared to their EGFR– counterpart, when grown under low-adherence neural stem cell condition medium (n=5 TLE samples, 10cells/μl (2000 cells/well); 1-3 wells per sample, week 2). Box-plots represent median, minimum and maximum value. P-value calculated using one-tailed U-Mann Whitney non-parametric test. **(g)** Representative microscope images (10X) of proliferative clusters growing from TLE EGFR+ cells. **(h)** Human TLE tissue immunofluorescence demonstrates occasional co-localization of the rare EGFR+ cells with the proliferative marker Ki67 (D). In (b) nuclei are counterstained with DAPI. Scale bar = 50μM unless otherwise specified.

Finally, to study functionally the proliferative properties of TLE glia, we employed an EGF-based purification strategy which has been used previously to isolate stem cell astrocytes from the adult rodent subventricular zone ^52^ and proliferative human stem cell populations from germinal matrix and glioblastoma fresh tissue samples ^53–55^. A quick analysis of the scRNA-seq data showed that EGFR expression was localized to a subset of cells from the following lineage clusters: astrocytes, OPCs, astrocyte/OPCs, pericytes, inflammatory cells and endothelial cells (Fig. 6d). Using EGF as a positive EGFR selection marker and CD34/45 to exclude endothelial and inflammatory cells, we detected a small but distinct population of EGFR+ (CD34–CD45–) cells (Fig. 6e) from five different pathological epilepsy tissues (Table 1), which we purified for downstream functional characterization of proliferation. Overall, EGFR+ cells formed significantly more proliferative clusters than EGFR– cells under serum-free, stem cell medium conditions (Fig. 6f-g), corroborating in vitro the proliferative phenotype previously observed in vivo (Fig. 6b, 6h).

## DISCUSSION

Accumulating scientific evidence in the past two decades has re-shaped the view of epileptogenesis as a neurocentric disease. In addition to aberrant synchronization in neuronal firing activity, several biological changes have been implicated in the pathophysiology of epilepsy, including gliosis, neuroinflammation, and reorganization of the ECM^6^. However, the specific molecular changes of each distinct neural cell type contributing to these pathological changes in epilepsy remain poorly understood^56^, especially in the primary human disease. To this end, here we developed a new immunotagging strategy to simultaneously isolate neuronal, astrocyte and OPC-enriched nuclei populations from bulk fresh-frozen human brain tissue, using antibodies against NEUN, PAX6, and OLIG2. This method allowed us, for the first time, to perform direct comparative analyses of the nuclear RNA-seq transcriptome in neurons, astrocytes and OPCs between control and diseased epileptic human temporal neocortex. Our analyses captured a rich repertoire of unique nuclear transcripts within each population at high throughput, and elucidated further the distinct transcriptome programs dysregulated in neurons, astrocytes, and OPCs in the context of medically refractory human TLE. We discovered an aberrant progenitor-like transcriptome phenotype in PAX6+ sorted epilepsy astrocytes, which we resolved further using independent single cell RNA-seq TLE dataset, confirmed in-situ both in human TLE and in a mouse epilepsy model, and found to be partially associated with upregulated EGFR expression and proliferation in-vitro.

Recently, several methods have emerged for isolating resting astrocytes and oligodendroglia acutely from human or mouse brain by means of immunopanning^27,31,44^. Others have used the surface marker GLAST to isolate astrocytes by FACS^57^, with some limitation on yield due to the sensitivity of GLAST+ cytoplasmic processes to mechanical dissociation and enzymatic digestion. Nuclei isolation from snap-frozen tissue circumvents the problem of cytoplasmic cell-cell processes dissociation and minimizes transcriptome alterations or artifacts that may be incurred during purification or cell culture^58^. Previous attempts to characterize glia using this method have been largely limited to the analysis of NEUN– (“non-neuronal”) populations, which are fundamentally heterogeneous and include endothelium, pericytes, smooth muscles, microglia and other inflammatory cells, in addition to astrocyte and oligodendroglial lineages^25,59–61^. Few studies have used OLIG2 or SOX10 to isolate oligodendroglial populations from the NEUN– fraction^62^ (unpublished), and a single study has demonstrated the use of SOX9 to sort astrocyte nuclei from mouse brain^27^. No studies thus far have demonstrated the simultaneous isolation of two defined glial lineages, astrocytes and OPCs, from the NEUN– fraction.

An important aspect to the isolation strategy presented in this study is its ability to distinctively enrich for oligodendroglial progenitors, rather than all oligodendroglial lineages, allowing isolation and comparative analysis of OPCs and astrocytes from the same tissue niche. For OPCs, we chose to sort using OLIG2 because this TF shows higher expression in OPCs compared to myelinating oligodendroglia, and has been used to distinguish the two populations by flow^62^. To ensure OPC enrichment, we excluded from our OPC gate the lower end of the OLIG2+ FANS population (i.e. OLIG2^LOW^), which we found to be enriched for myelinating oligodendroglia. Given the pivotal role of astrocytes and OPCs in development and the increasing appreciation for their contribution to neurological dysfunction, the simultaneous FANS isolation of these two distinct glial subtypes from banked fresh-frozen brain tissue is a valuable resource for further glial-specific omics analyses in the context of various pathological human CNS disorders. It also enables enrichment for astrocyte and/or OPC nuclei populations in single nuclei RNA-seq experiments, via hashing.

Previous bulk RNA-seq studies have begun to define the transcriptome in epilepsy mouse models and TLE patients^63,64^, but cell type-specific contributions have not been thoroughly characterized. Our study used two independent analyses to characterize the transcriptome of primary human TLE astrocytes, OPCs and neurons – FANS nuclei RNA-seq using fresh-frozen tissue and single cell RNA-seq using fresh tissue, providing a higher resolution of glial heterogeneity in primary human TLE neocortex. Differential expression analyses uncovered many of the biological changes previously implicated in epileptogenesis, and additionally resolved their cell type-specific transcriptome contribution. Neuronal TLE-dysregulated genes related to signal transduction, cell communication, and synaptic exocytosis, consistent with the model of epilepsy as a disease of imbalanced excitatory/inhibitory signaling^5,6,65,66^. TLE OPCs, rather, showed dysregulated functional pathways related to development, mitotic activity, cell-cell adhesion, TCA metabolism, and myelin sheathing, in line with their progenitor-like phenotype and tendency to increase in number at the site of brain injury, whether through local proliferation or via migration^7–13^. Given the complex biology of astrocytes and their still poorly understood functional and molecular heterogeneity in human disease^6^, we focused our main downstream analysis on the dysregulated gene pathways in abnormal TLE astrocytes isolated from their primary pathological niche. Comparative nuclear RNA-seq analysis in epilepsy (vs. control) astrocytes revealed robust downregulation of several astrocyte function-defining genes, including the transporters *SLC1A3*, *SLC1A2,* and *SLC1A4* and the gap junction proteins connexin 30(*GJB6*) and 43(*GJA1*). These findings are consistent with a large body of literature implicating astrocytic glutamate dysregulation in neuronal excitotoxicity and epilepsy^6,67–72^ although some primary human vs. experimental rodent TLE connexin expression differences were also noted^73^. Some of the most strongly and uniquely upregulated genes in epilepsy (reactive) astrocytes related to development, ECM/tissue repair, and potassium ion channels (*KCNK12*, *KCNK10, PKD2*), the latter two well documented in the dysfunction of astrocytes in the context of epilepsy^6,72,74^.

To increase the granularity of our glial cell type-specific transcriptome analysis, we performed single cell transcriptomics on temporal neocortex derived from five additional patients with medically refractory TLE. In this validation dataset, we were able to confirm the downregulation of canonical (mature) astrocyte markers and upregulation of developmental genes in TLE astrocytes. Moreover, bioinformatic analysis using three independent cell-clustering algorithms – Cell Ranger, Seurat, and MetaCell, uncovered a population of cells simultaneously co-expressing astrocytic and OPC markers. Importantly, such astrocyte/OPC cell cluster was not previously reported in the single cell RNA-seq analysis of healthy adult human temporal lobe tissue away from the seizure focus in patients with TLE^48^. Our findings are in accordance with the only other human epilepsy (population) transcriptome dataset, generated from HepaCAM-purified sclerotic hippocampus TLE astrocytes, which similarly found upregulation of OPC-like genes, such as *OLIG1, OLIG2, SOX10*, and *PDGFRA*, and downregulation of mature astrocytic markers, such as *GJA1* and *SLC39A12*, in hippocampus TLE compared to normal temporal cortex mature astrocytes^31^.

To confirm the existence of this hybrid astrocyte/OPC population in the context of epilepsy, we turned to a controlled mouse model of chronic seizure activity, where status epilepticus seizures are induced by pilocarpine. Indeed, glia co-expressing Gfap and Olig2 were noted only within the markedly gliotic neocortex of mice treated with pilocarpine, but not in controls, and this hybrid Gfap+/Olig2+ glia morphologically resembled reactive astrocytes. Few prior studies have noted the occasional expression of Olig2 by astrocytes under physiological conditions, in non-temporal brain regions^75–78^. Although we cannot completely rule out that the OPC-like reactive astrocytes discovered herein resulted from the expansion of pre-existent OLIG2-lineage astrocytes, previous electrophysiological analysis of sclerotic epileptic hippocampus has shown that TLE astrocytes can acquire a functional phenotype resembling NG2 glia^19^ suggesting a model of an aberrant phenotypic reversion of a subset of reactive astrocytes to a more immature, OPC-like state as a consequence of seizure activity. In line with the proposed immature phenotype, a subset of TLE astrocytes (astrocyte/OPCs) showed abnormal proliferation in-situ, and TLE glia expressing EGFR formed proliferative clusters in-vitro. Studies primarily in rodents have demonstrated that reactive astrocytes can revert into a developmental-like state after traumatic and ischemic injury, displaying aberrant proliferation and other stem cell characteristics^79–81^, and EGFR-mediated signaling appears to play an important role in the activation of injury-induced reactive astrocytes^51,82^. The causes and consequences of the aberrant immature phenotype in adult human astrocytes, discovered herein in the context of temporal lobe epilepsy, deserve further exploration in future studies.

## METHODS

### Sample collection

All tissue samples were obtained de-identified through the biorepository at the Icahn School of Medicine at Mount Sinai (ISMMS) under approved Institutional Review Board (IRB) protocols under consultation with the Ethics Committee of the Medical Board at the Mount Sinai Hospital. The study was performed under IRB-approved guidance and regulations to keep all patient information strictly de-identified, encrypted by the biorepository, and to only use excess tissue, not necessary for clinical diagnosis. For all subjects with potential use of identifiable information, informed consent was obtained prior to surgery, or from next of kin in the case of post-mortem tissue. IRB-approved waiver for consent was used for tissues that cannot be identified prospectively or retrospectively. No identifiable information or images are included in this study. De-identified epilepsy tissue was obtained within 5-30 minutes of surgical resection, from patients with medically refractory TLE with recent depth electrode recording of primary epileptic activity. De-identified control tissue was obtained from autopsy adult temporal lobe neocortex (TL) or pediatric germinal matrix with post-mortem interval less than 24 hours and without diagnostic neuropathological abnormalities. Tissue was either fresh-frozen for FANS, fixed in 4% paraformaldehyde for immunofluorescence studies, or collected fresh in live cell medium (PIPES) for single cell dissociation.

### Fluorescence Activated Nuclei Sorting (FANS)

We modified the existent FANS protocol for isolation of NEUN+ nuclei from fresh-frozen human brain cortex ^22,23^ by including positive selection for astrocyte (PAX6+ or SOX9+) and OPC (OLIG2+) enriched populations. Briefly, 200-500mg of frozen tissue was first manually homogenized using dounce glass grinders (Wheaton; 50 strokes) in a hypotonic lysis buffer (0.32M Sucrose/5mM CaCl_2_/3mM Mg(Ac)_2_/0.1mM EDTA, 10mM Tris-HCL pH8/1mM DTT/0.1% Triton X-100). Nuclei were collected in this buffer and purified from cellular debris by ultracentrifugation (107163.6 x g for 1h at 4°C) in a sucrose gradient (61.8%) (1.8M Sucrose/3mM Mg(Ac)_2_/1mM DTT/10mM Tris-HCL pH8). The nuclei pellet was resuspended in 1X PBS, and then nuclei were simultaneously incubated with the three fluorescently-conjugated primary antibodies (0.1% BSA/1X PBS) for one hour at 4°C on rotation: mouse anti-NEUN-AF555 (Millipore, MAB377A5, 1:1000); mouse anti-PAX6-APC (Novus Biologicals, NBP2-34705APC, 1:1000) or mouse anti-SOX9-AF647 (BD, 565493, 1:1000); and mouse anti-OLIG2-AF488 (Millipore, MABN50A4, 1:1000). DAPI (1:1000) was added after primary antibody incubation and before FANS (FACSAria™ III sorter; BD Biosciences). Nuclei were collected in Trizol LS (Life Technologies; 3:1 Trizol:nuclei ratio; for up to 50K nuclei) or in regular Trizol (Life Technologies; 750ul; for more than 50K nuclei), after first concentrating the solution in 1XPBS containing 0.36M Sucrose, 3.6mM Mg(Ac)_2_, 2mM Tris-HCL pH8, 5mM CaCl_2_, and were snap-frozen at −80°C for subsequent RNA isolation.

### Bulk RNA-seq preparation and analysis

Total nuclear RNA was isolated from FANS populations (NEUN+, NEUN–OLIG2+ and NEUN–PAX6+) by standard phenol /chloroform extraction, followed by DNase digestion on tube (15 min), and RNA cleanup and concentration in final volume of 15μL water (Zymo Research, R1013). RNA concentration was determined using Qubit (ThermoFisher). For FANS RNA-seq library preparation (SMARTer Stranded Total RNA-Seq Kit Pico Input Mammalian, Clontech Laboratories, 635005), 2952pg of total nuclear RNA were amplified into cDNA with fragmentation times of 2.5 min for autopsy and 3.5 min for TLE cases (14 cycles total amplification for all). Given the postmortem nature of TL specimens, we employed cDNA synthesis and library preparation kit optimized for partially degraded RNA with simultaneous depletion of ribosomal RNA, which passed quality control requirements. Ribosomal RNA depletion was performed using human-specific R-Probes. Libraries were generated using Nextera XT (FC-131-1024) and validated using Agilent 2100 Bioanalyzer. Sequencing was performed on Illumina HiSeq 2500 (50bp pair-end sequencing, 38–50 million paired-end reads/sample).

Sequenced output FASTQ files of bulk RNA-seq data were assessed for quality using the FASTQC package. Reads were aligned to the human genome (GENCODE GRCh38) using STAR with default settings ^83^. Gene counts were obtained using the featureCount utility ^84^. The counts data were rld (rlog transformed counts)-normalized. Differential expression analysis was performed using the DESeq2 R package ^85^, modeling the data with a negative binomial distribution and using Empirical Bayes shrinkage for dispersion and fold change estimation. Functional and gene set enrichment analyses were performed using several tools: HOMER ^86^ with background defined as the set of genes passing the independent filtering low expression threshold by DESeq2; DAVID ^87^, and GSEA ^43^.

### RT-qPCR analysis

FANS RNA was also used to generate cDNA for RT-qPCR (High-Capacity RNA-to-cDNA Kit, Life Technologies, 4387406). Real time PCR was run in duplicates using the SYBR-Green system (Quanta Biosciences, 101414) (7900HT, Life Technologies). Primers spanned exon-exon junctions; melting curves were analyzed to ensure primers specificity; genomic DNA was used as negative control. Fold gene expression was calculated as 2^−(CtSample – CtHKG)^ relative to the NEUN+ fraction (2^−ΔΔCt^). ACTB was used as housekeeping gene, HKG.

### Droplet-based single cell RNA-seq preparation and analysis

TLE tissue from five different de-identified patients was obtained from the operating room and immersed in freshly prepared live cell buffer (PIPES) for single cell dissociation as described for FACS ^54^. Single cell RNA-seq was performed on these samples using the Chromium platform (10x Genomics, Pleasanton, CA) with the 3’ gene expression (3’ GEX) V1 kit for 2 samples, V2 kit for 2 samples, and V3 for 1 sample, using an input of ~10,000 single cells. Briefly, Gel-Bead in Emulsions (GEMs) were generated on the sample chip in the Chromium controller. Barcoded cDNA was extracted from the GEMs by Post-GEM RT-cleanup and amplified for 12 cycles. Amplified cDNA was fragmented and subjected to end-repair, poly A-tailing, adapter ligation, and 10X-specific sample indexing following the manufacturer’s protocol. Libraries were quantified using Bioanalyzer (Agilent) and Qubit (Thermo Fisher) analysis. Libraries from 1000-4000 cells (depending on the sample) were sequenced in paired end mode on a HiSeq 4000 instrument (Illumina, San Diego, CA) at an average depth of 174,446 reads per cell (for V2/V3). For analysis, the Cell Ranger v3 software was first used to demultiplex cellular barcodes to produce raw 3’ end read profiles for individual cells, and then performed alignment, filtering, barcode counting, and unique molecular identifier (UMI) counting, producing a feature-barcode matrix per sample. Unsupervised clustering analysis was performed by three independent methods: Cell Ranger (10X Genomics), Seurat v2.3.4 4 ^45^, and MetaCell ^46^. Doublet rate was estimated at 1-5% using a linear approximation of ~4.6e-06 * [#_of_cells_loaded] (https://satijalab.org/costpercell) and considered further during the data analysis based on non-random distribution of UMI cell counts. Batch correction and integration of cell populations between samples was done using mnnCorrect ^47^. For Seurat analysis, the final data matrix was produced by removing unwanted cells (with too few UMIs, set at default value, or cell with overall expression attributable to mitochondrial genes exceeding 5%) from the dataset, and applying a global-scaling normalization method that normalizes the feature expression measurements for each cell by the total expression, multiplies this by a scale factor (10,000 by default), and log-transforms the result. For MetaCell analysis, the raw data matrix was filtered to remove specific mitochondrial genes, and then cells with less than 500 UMIs. MetaCell performs a cell-to-cell similarity matrix normalization in the process of constructing its K-nn graph, enforcing it to have a more symmetric structure, with more balanced in-going and out-going degrees.

### Pilocarpine-induced epilepsy mouse model

Animal studies were performed in accordance with the ethical standards at the Icahn School of Medicine at Mount Sinai (ISMMS) under an approved Institutional Animal Care and Use Committee (IACUC) protocol. Male C57BL/6NCrL mice (Charles River, France) weighing 19–22 g (7 weeks old) were used. Seizures were induced through single pilocarpine injection intraperitoneally (ip, 285 mg/kg) 30 min after scopolamine methyl bromide (1 mg/kg, ip) administration. Behavioral observations were performed, and seizures were video recorded until mice entered into status epilepticus (SE) (Racine scale level 5 tonic clonic seizures/continuous shacking). Mice developing SE were injected with a dose of diazepam (10 mg/kg; ip) after 2h to limit the duration of SE and monitored to ensure cessation of convulsive behavior. Saline solution was injected (1cc, ip) to compensate for body fluid lost during SE. Control mice were injected with equal volume of saline solution at the time of scopolamine injection. Mice were euthanized seven weeks after the onset of SE (650 mg/kg pentobarbital, ip) and transcardially perfused (1X PBS followed by 4% PFA/1X PBS). Brains were post-fixed in 4% PFA/1X PBS overnight, cryoprotected through gradual immersion in 12.5 to 25% sucrose gradient for 48h, embedded in O.C.T, and serially sectioned using a cryostat (25μM thickness).

### In vitro proliferation assay

To assess proliferation, EGFR+ (CD34–CD45–) and EGFR– (CD34–CD45–) cells were isolated from ~500mg of fresh human TLE tissue as previously described ^53,54^, and were seeded immediately after FACS on 96-well low-adherence plates at a density of 10c/μl, in triplicates, in NS media (1X N2, 1X B27, 20μM glutamine, 1X Insulin/Transferrin/Selenium, 15mM HEPES, 0.6% glucose, 1X Antibiotic/Antimycotic, in DMEM/F12 media) supplemented with EGF (20ng/ml) and bFGF (20ng/ml). Cells were maintained at 37°C and 5% CO2 changing 2/3 of media on day 6 and every 3-4 days thereafter. Images of NS formation were captured with a light inverted microscope (Motic AE31) 2 weeks after seeding. Pictures covering the entire surface of the wells were taken at 10X and were used for subsequent counting.

### Immunofluorescence

Specimens were fixed in 4% paraformaldehyde/1X PBS at 4°C for 24h (TL/TLE cases), and up to 72h in the case of germinal matrix, rinsed in 1X PBS, and vibratome sectioned (30 μm). Sections were incubated for 1 hour in blocking solution (1X PBS/0.5% Triton X-100/10% normal donkey serum); then for 24h at 4°C in primary antibody (1X PBS/0.25% Triton X-100/1% normal donkey serum); and then for 4 hours at room temperature in either donkey anti-mouse, donkey anti-rabbit, donkey anti-rat, or donkey anti-goat fluorochrome-conjugated secondary antibodies (Jackson Laboratories, 1:250 dilution). Formalin-fixed paraffin embedded (FFPE) tissues underwent 1hr deparaffinization, rehydration in decreasing gradient of ethanol, and antigen retrieval for 20min prior to blocking.

Primary antibodies dilutions were as follows: 1:50 mouse anti-EGFR (Invitrogen 280005); 1:500 rat anti-GFAP (Life Technologies, 13-0300); 1:250 rabbit anti-OLIG2 (Millipore, AB9610); 1:250 goat anti-hOLIG2 (R&D Systems, AF2418SP); 1:250 rabbit anti-Ki67 (Abcam, Ab15580); 1:250 mouse anti-Ki67 (BD Biosciences, 556003); 1:100 rabbit anti-PAX6 (Novus Biologicals, NBP1-89100); 1:100 mouse anti-NEUN (Millipore, MAB377); 1:250 rabbit anti-AIF1 (IBA1) (Wako, 019-19741). Nuclei were counterstained with DAPI (1:1000). Images were obtained using a LSM 780 upright confocal microscope (Zeiss).

### Statistical analysis

Two-tailed and one-tailed unpaired Students’ t-test was used to calculate significance (*p<0.05, **p<0.01, ***p<0.001), assuming homogeneous variances. For non-parametric analysis, we used the Mann-Whitney U test. Bar graph data is represented as mean ± SEM of at least three independent experiments. All RNA-seq tests were FDR adjusted for multiple testing correction.

## ETHICS APPROVAL AND CONSENT TO PARTICIPATE

All tissue samples were obtained de-identified under approved Institutional Review Board (IRB) protocols and appropriate consent. Animal studies were performed in accordance with the ethical standards at the Icahn School of Medicine at Mount Sinai (ISMMS) under an approved Institutional Animal Care and Use Committee (IACUC) protocol.

## CONSENT FOR PUBLICATION

All authors have approved the manuscript for publication.

## AVAILABILITY OF DATA AND MATERIAL

The data that support the findings in this study are publicly available at https://data.mendeley.com/drafts/w4d7sdc629, and are currently being deposited in NCBI’s Gene Expression Omnibus to be accessible through a GEO Series accession number GSE140393 prior to publication.

## COMPETING OF INTERESTS

The authors declare that there is no conflict of interest regarding the publication of this article.

## FUNDING

The research was supported by R03NS101581 and RF1DA048810 (N.M.T.) and R01MH110555 (D.P.) NIH funding.

## AUTHOR CONTRIBUTIONS

Conception of project: N.M.T. Experimental design and data interpretation: N.M.T., E.Z., J.T-G., and G.N. Tissue procurement: S.G., F.P., K.S., L.M., J.Y., D.P. Development and implementation of FANS strategy: J.T-G., Y.J., S.A, N.M.T. Nuclear and single-cell RNA-seq preparation: J.T-G., K.A., K.G.B., R.S. Bioinformatic analysis: G.N., E.Z., Y.W., K.G.B. FACS, cell culture, and histology: J.T-G., E.C. Pilocarpine mouse model: M.D., A.S. Manuscript preparation: N.M.T. and J.T-G., and editing: all authors.

## ACKNOWLEDGEMENTS

We thank members of the Pathology and Neurosurgery departments and the Biorepository Core at ISMMS for facilitating collection and procurement of de-identified tissue under appropriate consent; the ISMMS Flow Cytometry Core Facility for accommodation of human tissue sorts 24 h/day; the ISMMS Genomics Core Facility for computing resources and sequencing; and the NYGC for sequencing.

## Notes

### Competing Interest Statement

The authors have declared no competing interest.

