## Supplemental information for "Cell type-specific isolation and transcriptomic profiling informs glial pathology in human temporal lobe epilepsy"

Jessica Tome-Garcia^1,2^, German Nudelman^3^, Kristin G. Beaumont^4,5^, Ying-Chih Wang^4,5^, Mary Duff^2^, Saadi Ghatan^6^, Fedor Panov^6^, Elodia Caballero^1,2^, Kwadwo Sarpong^4,5,7^, Kimaada Allette^4^, Lara Marcuse^3^, Jiyeoun Yoo^3^, Yan Jiang^2,7^, Anne Schaefer^2^, Schahram Akbarian^2,7^, Robert Sebra^4,5^, Dalila Pinto^4,5,7^, Elena Zaslavsky^3^, and Nadejda M. Tsankova^1,2*^

**SUPPLEMENTARY INFORMATION**

**Supplementary Fig. 1. Immunotagging strategy for simultaneous isolation of astrocyte, neuronal, and OPC-enriched populations from human temporal lobe neocortex. (a)** Representative FANS pseudocolor plots showing sequential gating of non-debris (red), non-doublets (green and blue), live DAPI+ nuclei (purple), NEUN+ neuronal (red), PAX6+( NEUN–) astrocyte (orange), OLIG2+( NEUN–) OPC-enriched (green), and OLIG2^LOW^ (purple) mature oligodendroglial-enriched nuclei populations from postmortem TL control fresh-frozen neocortex. TN = triple negative (PAX6–OLIG2– NEUN–) population (black). **(b)** FANS pseudocolor plots showing alternative astrocyte isolation strategy of astrocytes from TL neocortex using SOX9 instead of PAX6, in combination with NEUN and OLIG2. **(c)** Expression of *GFAP* and *CD44* assessed by RT-qPCR in SOX9+, PAX6+ and other FANS populations derived from TL tissue. **(d)** Expression of the microglial marker *CD11B (ITGAM)* assessed by RT-qPCR in TLE tissue. **(e)** Representative immunofluorescence images of NEUN, OLIG2 and GFAP expression in TL control tissue. Scale bar = 50µM.

**Supplementary Fig. 2. Hierarchical clustering of** **FANS populations according to cell type**. Unsupervised hierarchical clustering dendrogram and heatmap of all FANS nuclear RNA-seq datasets, using Euclidean distance as the metric, shows overall clustering of populations according to cell type, with some heterogeneity seen in PAX6+ TLE astrocytes.

**Supplementary Fig. 3. Single cell RNA-seq data from five TLE patients informs the presence of an astrocyte/OPC hybrid population.** T-distributed stochastic neighbor embedding (t-SNE) visualization of each individual TLE sample analyzed, K-Means clustering using Cell Ranger V3.0. Canonical cell type-specific markers are used to annotate the lineage of each cluster for each sample. More than one astrocyte cluster is detected in all but one of the samples, with an aberrant hybrid astrocyte/OPC cluster (underlined). Data is visualized using the Loupe Cell Browser, version 3.0.1.

**Supplementary Fig. 4. Distribution of canonical lineage markers used for single cell RNA-seq cluster annotation** Violin plots show Seurat cluster distribution and expression of selected canonical lineage markers for mature oligodendrocytes, neurons, lymphocytes, pericytes, smooth muscle cells (SMCs), microglia, and endothelial cells in samples 19619 (top) and 14431 (bottom). Predicted annotation of cell type for each cluster is indicated in each pertinent legend.

**Supplementary Fig. 5. MetaCell analysis disambiguates reactive astrocytes from OPCs (a)** MetaCell unsupervised clustering analysis distinguishes presumed reactive astrocytes from OPCs in sample 14431 based on expression of *GFAP* and other cell lineage markers (right) but fails to do so when using canonical astrocyte markers defined through normal surgical temporal neocortex single cell RNA-seq analysis ^48^ (left). **(b)** Heatmap of sample 14431 showing top highly expressed genes within each cell cluster identified by MetaCell. *GFAP*+ astrocyte (brown) and OPC (green) clusters are similar but distinct. (**c**) Functional enrichment analysis of differentially overexpressed genes in cells expressing high levels of *GFAP* (*GFAP*++ vs. everything else, > 2.5 log_2_ fold change) within the astrocyte/OPC cluster shows significant enrichment of processes related to astrocyte function and CNS development (p-value < 0.1).

**Supplementary Table 1.** Normalized RNA-seq gene expression data for all sorted nuclei, log-transformed (rld-normalized).

**Supplementary Table 2.** Differential expression, TLE vs. TL, nuclear RNA-seq datasets for astrocyte, OPC, and neuronal FANS populations, with annotation of dysregulated genes unique to each cell type.

**Supplementary Table 3.** Functional enrichment analyses of differentially expressed genes for each nuclei cell type, TLE vs. TL, and for the scRNA-seq analysis of 14431 astrocyte/OPC hybrid cluster vs. everything else, generated using DAVID.

**Supplementary Table 4.** Functional enrichment analyses of differentially expressed genes for each nuclei cell type, TLE vs. TL, generated using HOMER.

**Supplementary Table 5.** Sequencing and mapping quality control metrics of single cell RNA-seq data from five independent TLE samples. Samples 14431 and 19619 were of highest quality and best metric data and were used in further Seurat and Metacell analyses. List of top cluster-defining genes in Seurat and MetaCell analyses for samples 19619 (Seurat only) and 14431 (Seurat and MetaCell).

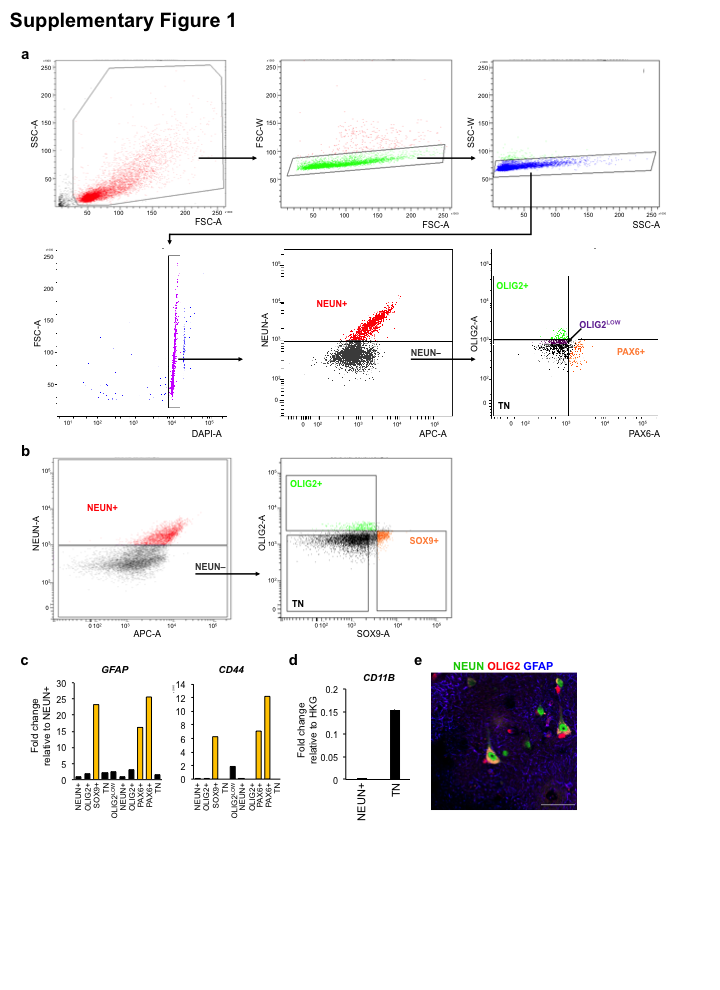

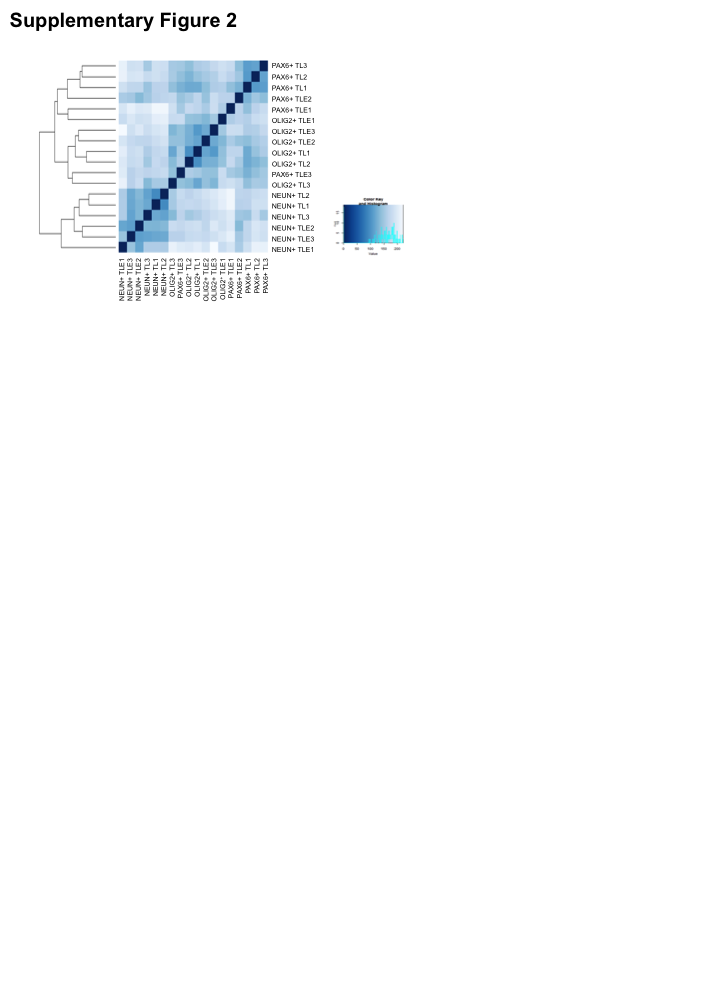

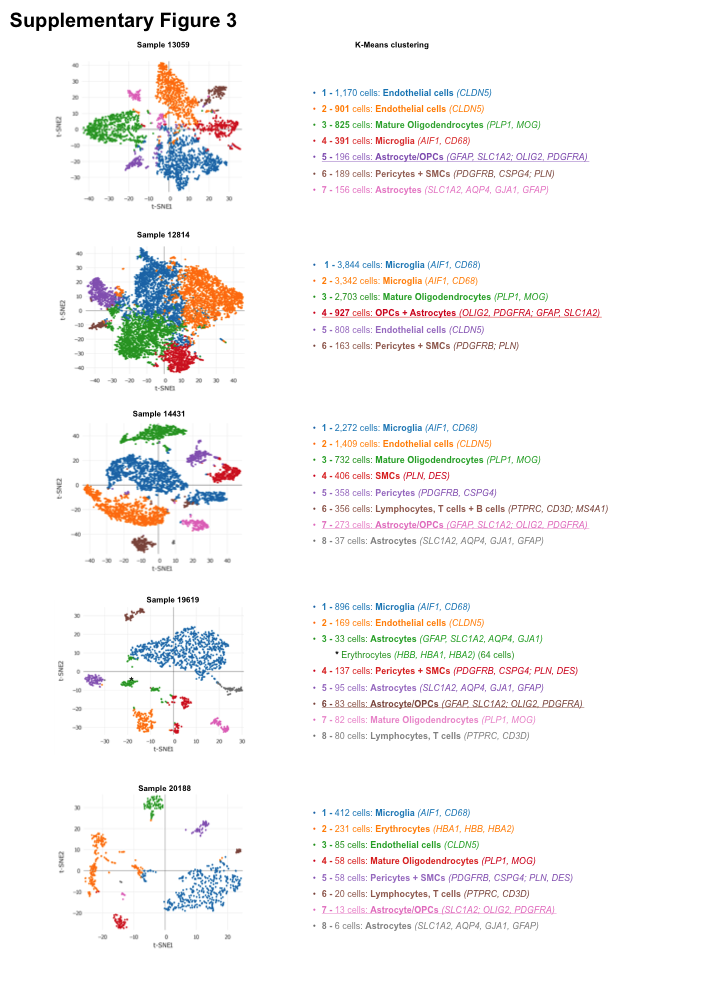

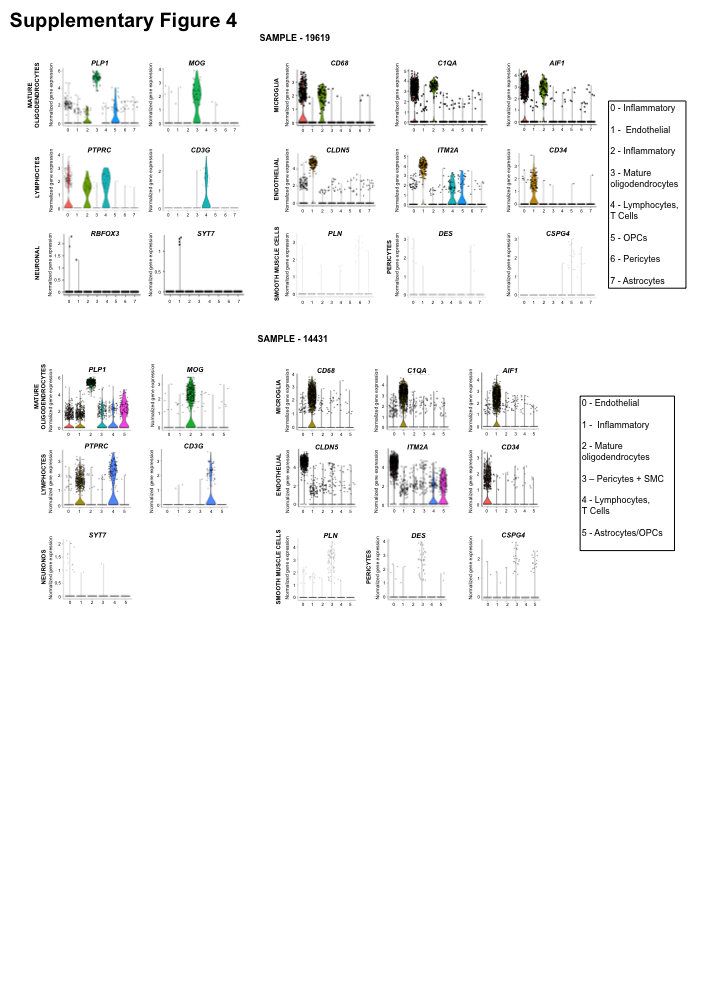

**Supplementary Figure 5**

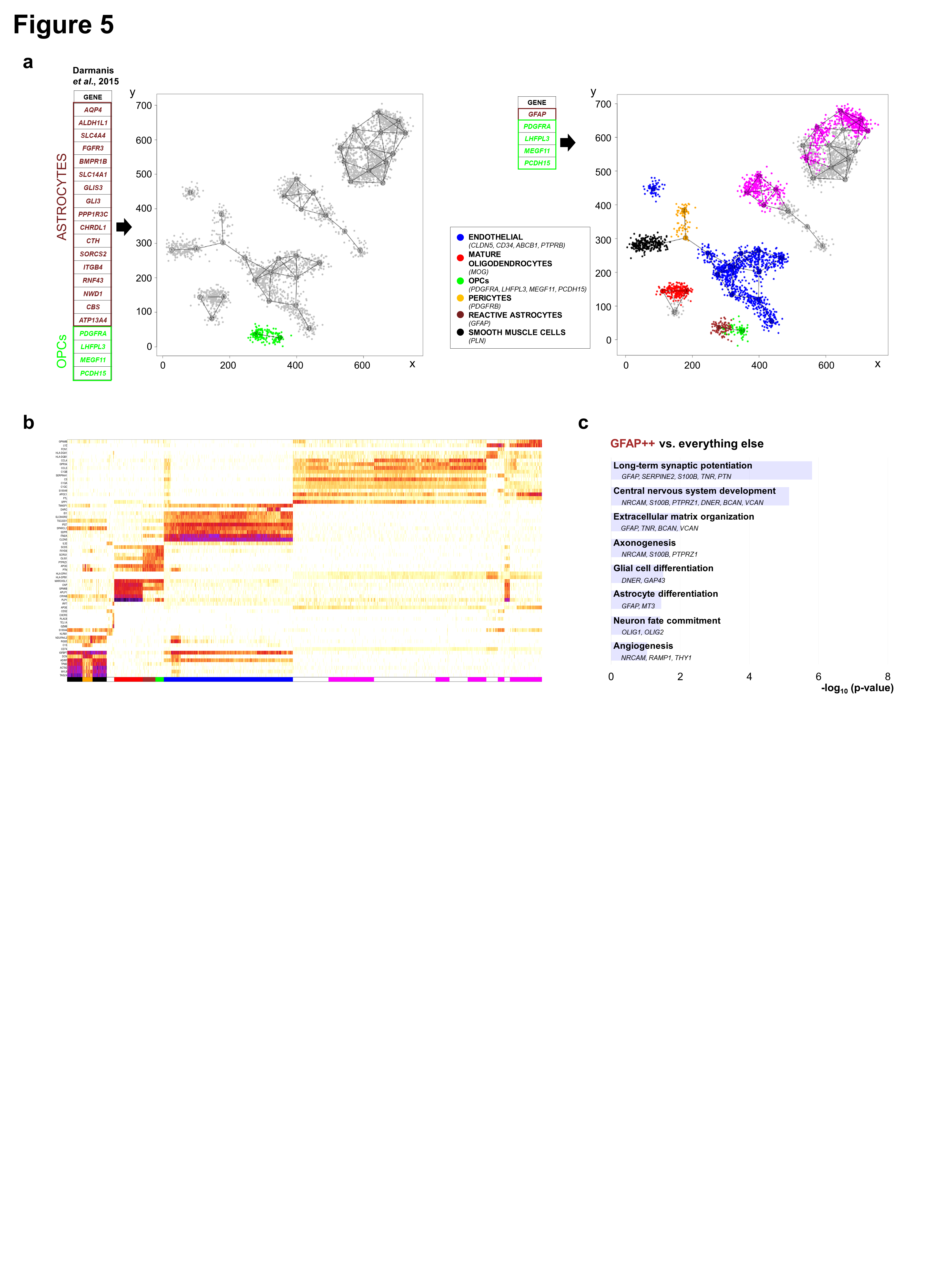
