## Supplementary figures and images for "Cell type-specific isolation and transcriptomic profiling informs glial pathology in human temporal lobe epilepsy"

### Supplemental Figure1

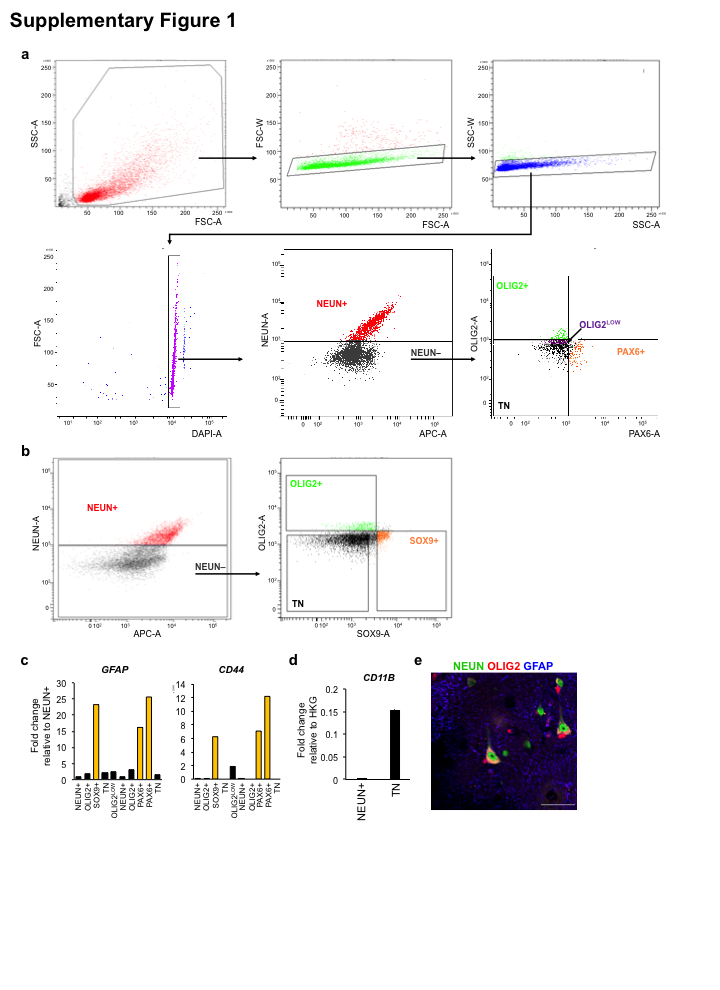

### Supplemental Figure2

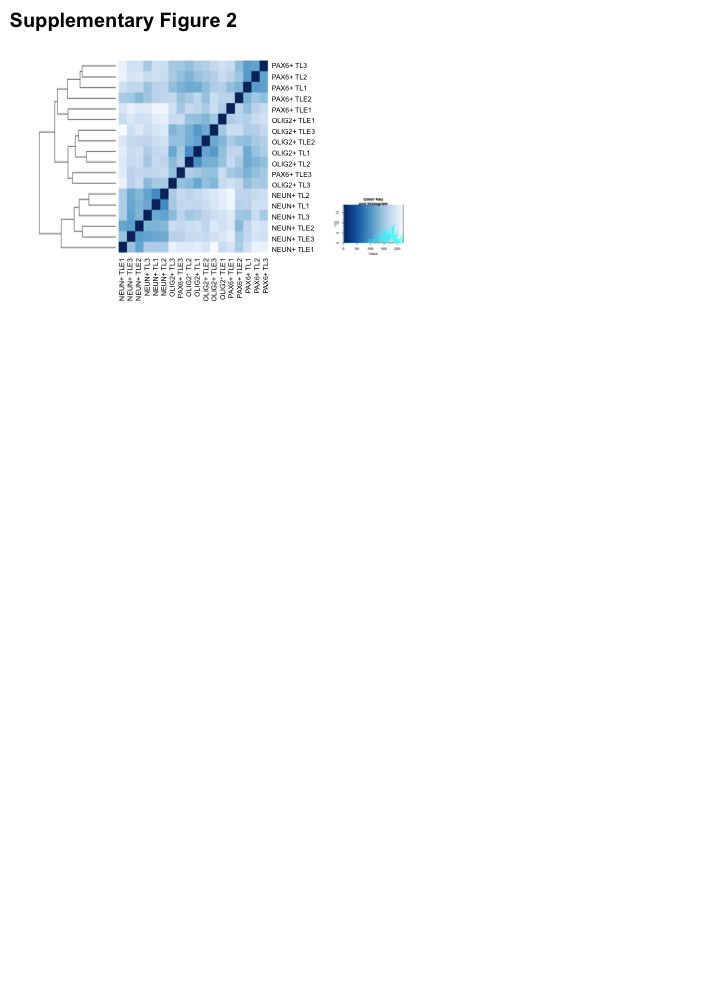

### Supplemental Figure3

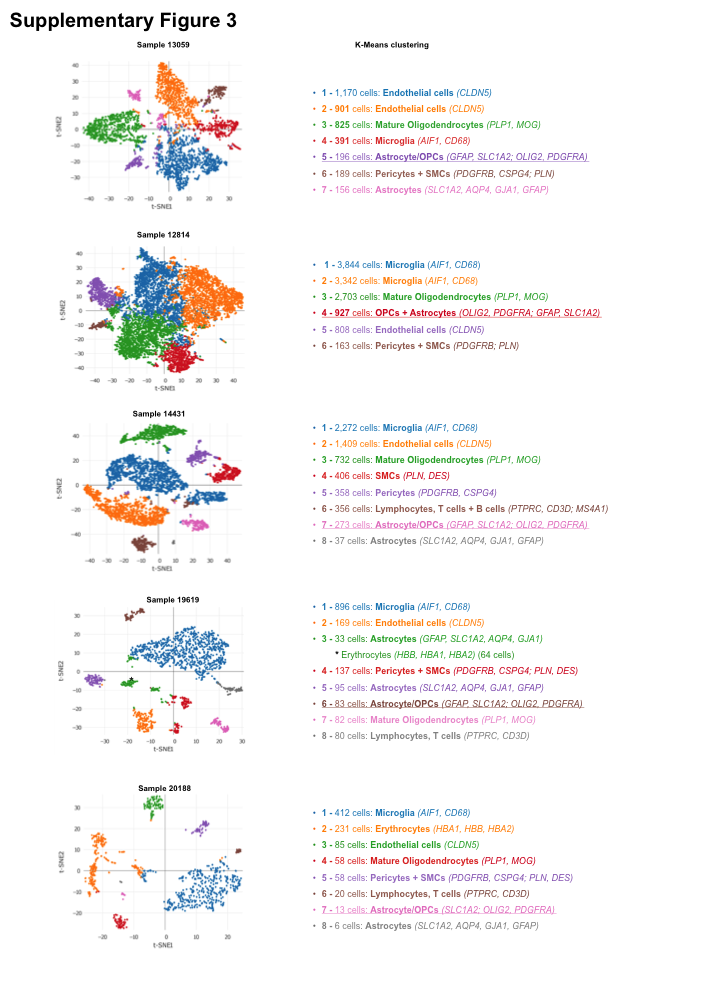

### Supplemental Figure4

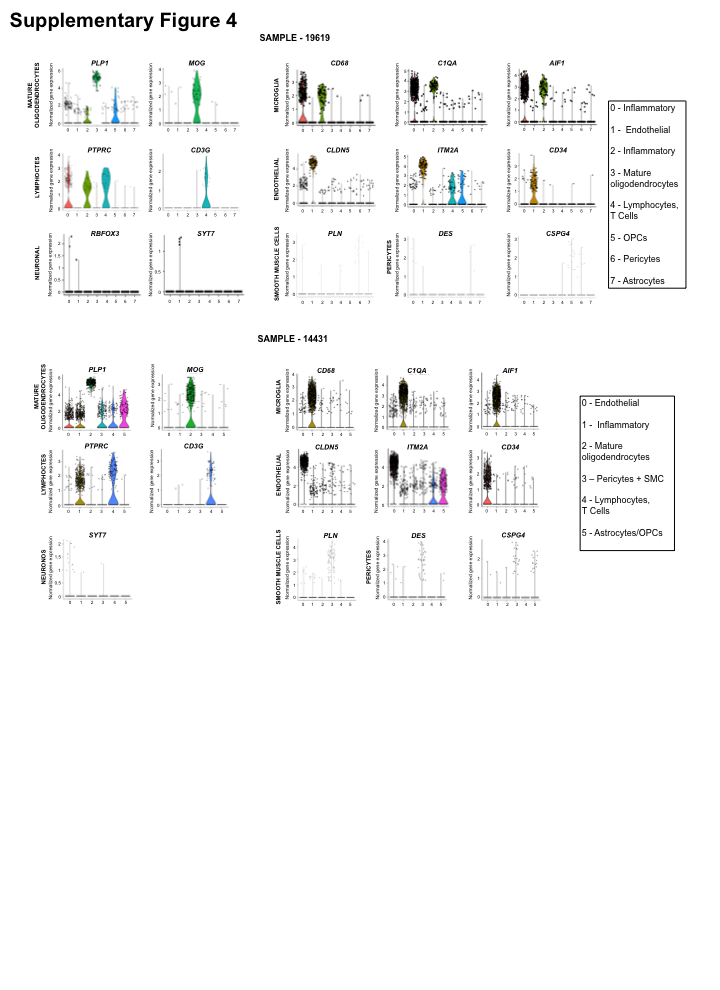

### Supplemental Figure5

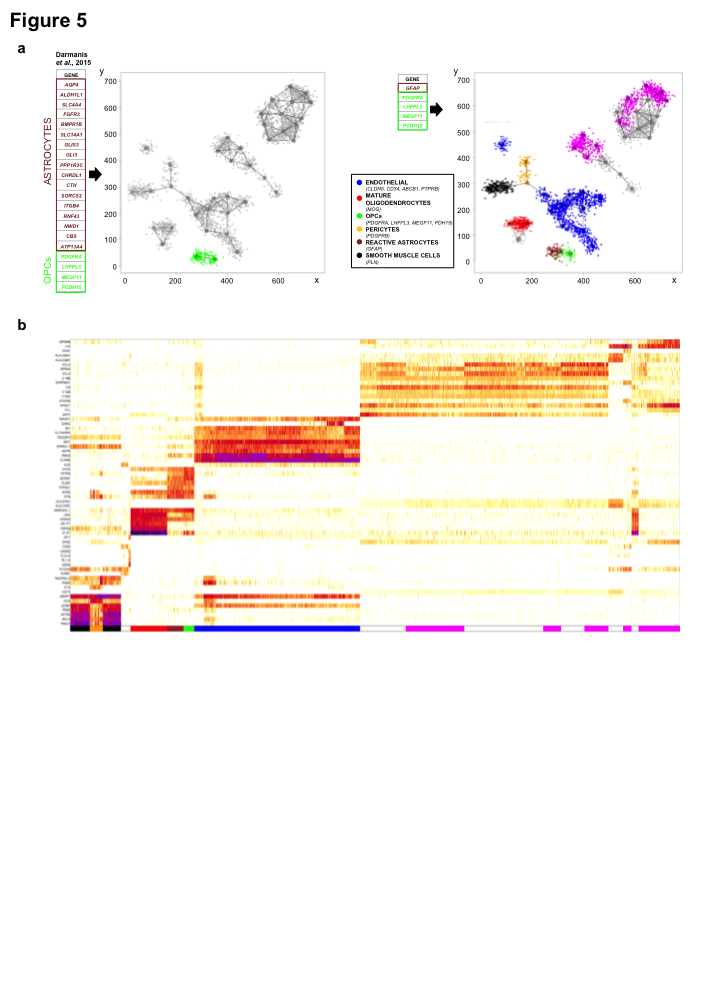
